# *Plasmodiophora brassicae* in Mexico, from anecdote to fact

**DOI:** 10.1101/2021.11.29.469274

**Authors:** Legnara Padrón-Rodríguez, Carlos Roberto Cerdán Cabrera, Nadia Guadalupe Sánchez Coello, Mauricio Luna-Rodríguez, Edel Pérez-López

**Affiliations:** Facultad de Ciencias Agrícolas, Universidad Veracruzana. Circuito Gonzalo Aguirre Beltrán s/n, Zona Universitaria, Xalapa, Veracruz, México; Centre de recherche et d’innovation sur les végétaux (CRIV), Université Laval, Department of Plant Sciences, FSAA, Université Laval

**Keywords:** Clubroot, cruciferous crops, broccoli, Puebla and Tlaxcala, ‘hernia de la col’

## Abstract

For years, the presence of clubroot disease and its causal agent, *Plasmodiophora brassicae*, in Mexico has been given by granted. However, after a long search in the scientific literature in English and Spanish, as well as grey literature including thesis and government reports, we were not able to find any information regarding the actual detection of the pathogen, hosts affected, areas with the disease, or any real information about clubroot (‘hernia de la col’, in Mexico). To confirm if *P. brassicae* was indeed in Mexico, we started a true detective adventure. First, we identified agricultural communities in south-east Mexico known to grow cruciferous crops. Second, we asked to the growers if they have ever seen clubroot symptoms, showing them during the inquires pictures of the characteristic galls that might have been present in their crops. Third, we collected soil from two of the communities with positive response and grew an array of cruciferous in the soil as baits to “fish” the clubroot pathogen. We detected the presence of galls in the roots of 32 plants and observed the presence of resting spores. Through a *P. brassicae* specific PCR assay, we were able to confirm the presence of the clubroot pathogen in the samples and in Mexico for the very first time. This study is the first report and identification of *P. brassicae* in Mexico, opening the doors to understand the genetic diversity of this elusive and devastating plant pathogen.

## INTRODUCTION

Clubroot is a devastating disease affecting cruciferous crops worldwide (Botero et al. 2019). Clubroot is caused by the obligate biotrophic, parasitic protist *Plasmodiophora brassicae* Woronin, 1877 (Burki et al. 2010). *P. brassicae* colonization and establishment in the susceptible host is divided in primary infection of a susceptible host plant, characterized by the intrusion of root hairs primary by zoospores, leading to the production of secondary zoospores that, during secondary infection of cortical cells can establish a nutrient-sink galls to support the production of new resting spores, remaining as a viable soil-borne pathogen for many years (Kageyama and Asano 2009).

The clubroot pathogen was first described in Russia (Woronin, 1877), although previous reports of the disease are as far dated as 1500s (Howard et al. 2010). Not long after, the clubroot disease started to spread through eastern and northern Europe, finding its way to The Americas, specifically Canada around the 1800s, maybe through the large influx of immigrants from eastern Europe (Minogue 2013). However, how the clubroot pathogen arrived in Latin America is still a mystery. Other theory places the point of entry of the clubroot pathogen to The Americas through Latin America. A report of clubroot-like symptoms on cabbage was made in Spain by Díaz de Isla in 1539 (Botero et al. 2019), indicating that the clubroot pathogen was probably present in Spain when Columbus “discovered” The Americas and could have been introduced during the subsequent colonization.

Unlike Europe and North America, most research on clubroot (*hernia de la colza* or *hernia de la col*, in Spanish) in Latin America has focused on cruciferous vegetables (García et al. 2006). The disease has been reported in South America affecting cabbage, Chinese cabbage, broccoli, arugula, cauliflower, and canola, and in Central America affecting broccoli and cabbage (Botero et al. 2019). However, the severity and economic losses caused by *P. brassicae* in the region are not known. In North America, mainly Canada, clubroot causes huge economic loses every year to the canola industry, reason why the focus has been on that crop, while in United States more and more canola producing areas are found to be affected by the disease every year (Northern Canola Growers Association, 2020).

The scenario in Mexico, another North American country, is different. Although it is mentioned in the European and Mediterranean Plant Protection Organization database (EPPO, 2019) and two published reviews (Dixon 2009; Botero et al. 2019) that clubroot is present in the country, we haven’t been able to find any study characterizing the pathogen in English or Spanish. Taking into consideration that Mexico is the world’s fifth largest broccoli producer, and the main provider of this vegetable to East United States and Canada, it is important to finally confirm the presence of not of the clubroot pathogen to be ready for potential outbreaks. To that end, we developed a real phytopathological detective work. First, we visited in Puebla and Tlaxcala, two of the main broccoli producer states looking for producers that might have seen clubroot symptoms in their fields. Later, using soil collected from those fields, we were able to identify clubroot-related symptoms in four different brassicas crops grown in them, confirming the presence of *P. brassicae* through molecular assays. This study is the first report of clubroot disease in Mexico and is the first step in our task of studying the world diversity of *P. brassicae*.

## MATERIALS AND METHODS

### Identification of study sites and soil collection

To identify the best suitable sites for soil collection, we focused on two of the main broccoli producer states in Mexico, Tlaxcala and Puebla. In Tlaxcala we interviewed several leaders of grower’ associations and in Puebla members of the local government, allowing the identification of fourteen fields suitable to collect soil samples that could have *P. brassicae* or that could have been affected in the past by the disease. The fourteen fields were located in two municipalities, Lázaro Cárdenas (LC), Tlaxcala, and San Pedro Cholula (SPC), Puebla, and grouped into three categories: (1) fields in production (LC6; 19°31’7.33”N, 97°59’8.56”W, SPC1; 19°5’10.26”N, 98°20’51.40”W, SPC3; 19°5’13.60”N, 98°20’52.35”W, SPC4; 19°3’9.42”N, 98°20’52.41”W, and SPC7; 19° 3’14.01”N, 98°20’50.67”W), (2) fields without brassicas from 2 months to 1 year (LC3; 19°31’51.76”N, 97°59’2.31”W, LC5; 19°31’50.21”N, 97°58’59.45”W, SPC2; 19°5’5.02”N, 98°20’55.27”W, SPC5; 19°3’8.43”N, 98°20’53.05”W, and SPC6; 19° 3’10.55”N, 98°20’50.04”W), and (3) fields without brassicas for 2 years or more (LC1; 19°31’53.14”N, 97°59’3.94”W, LC2; 19°31’55.18”N, 97°59’4.35”W, LC4; 19°31’47.60”N, 97°59’7.25”W, and LC7; 19°32’58.84”N, 97°59’8.88”W) (Fig. 1A).

**Fig. 1.**
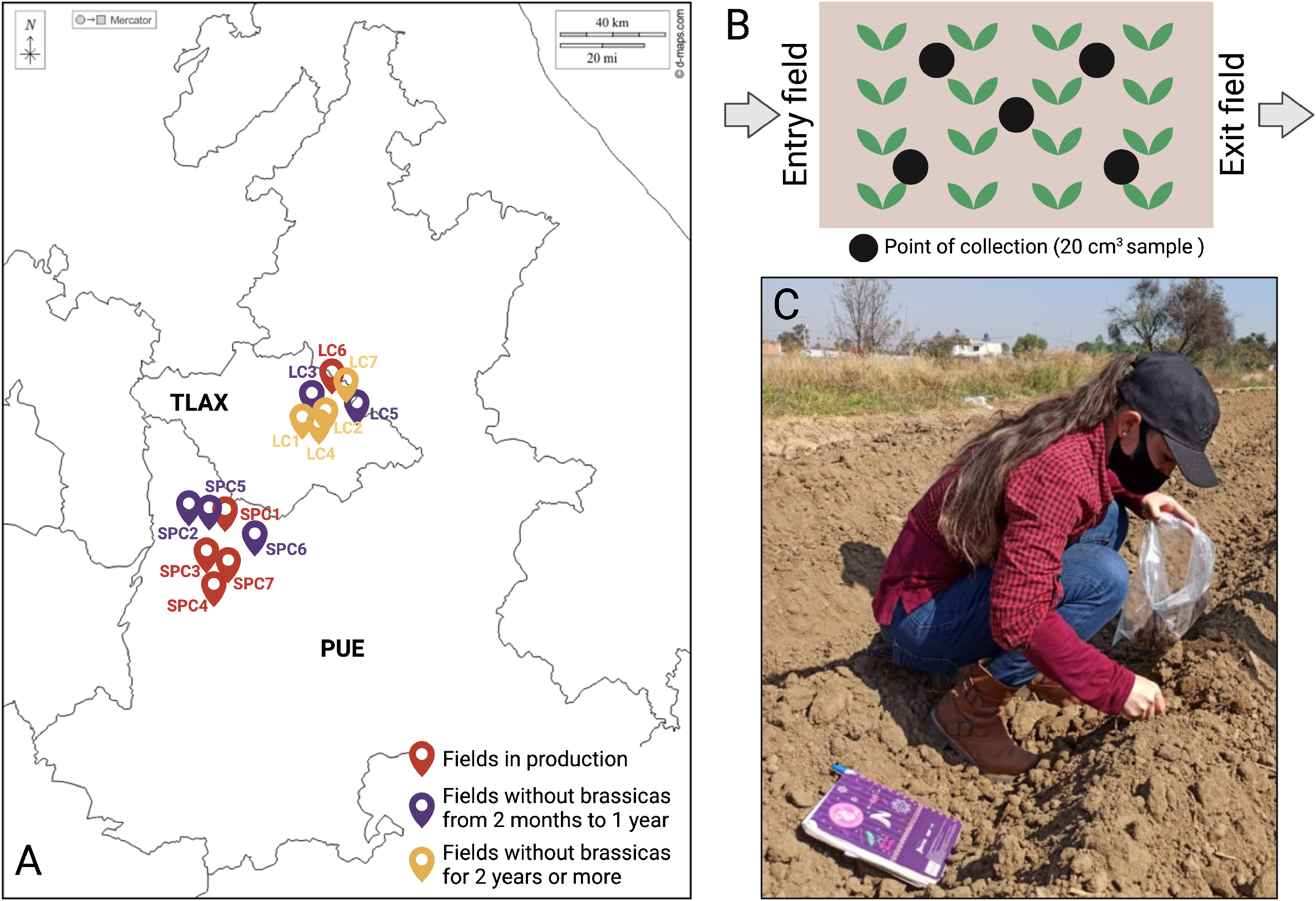
Soil collection in Puebla and Tlaxcala, Mexico. **A**. Map with the location of the 14 sites sampled in both states. In three different colors have been represented the 3 group of fields samples. **B**. Schematic representation of the field and the design in “M” performed covering all the field. **C**. Legnara during the soil collection in Tlaxcala, Mexico.

### Sample collection

Although several fields sampled were in production, not clear clubroot symptomatology such as yellowing or dehydration of the plant was detected in the foliage. That is the reason why we collected from all the fields only soil. Soil sampling was conducted following an “M” pattern that allowed to cover all the field and have samples from the entry point, the center, and the exit of the field (Fig. 1B-C). Each sample consisted of 20 cm^3^ of soil, including organic matter that could have been in the field. The samples were placed in plastic bags and translated to the laboratory. Seventy samples were collected. The five samples from each field were well mixed and after they were divided in two to proceed to “fish” *P. brassicae* resting spores, leaving us with 28 containers of soil. To avoid cross-contamination, shoes and tools were disinfected from one field to the other using bleach 15% solution.

### Plant material and infection assays

To “fish” *P. brassicae* resting spores, we used cabbage (*Brassica oleracea* var. *capitata* L.), broccoli (*Brassica oleracea* var. *italica* Plenck), cauliflower (*Brassica oleracea* var. *botrytis* L.), and long red radish (*Raphanus sativus* var. *sativus* L.) as bait. The seeds obtained from the distributor *Rancho Los Molinos* (Mor., Mexico) were disinfected and sown into semi-solid media MS (Sigma, USA). Ten days after germination, sixteen plants, four of each host, were transplanted into the soil collected from the fields in a completely randomized design. As negative control, we included one container with commercial peat moss-based substrate with similar characteristics to Sunshine Mix #4, which was incubated with 16 plants, establishing an overall of 480 interactions. Plants were grown under a 16/8-h light/dark cycle, 100 μmol (photons)/m^2^/s, and a constant 20 °C.

### *Plasmodiophora brassicae* detection

All plants were evaluated 60 days post transplantation (60 dpt) to soil collected from the fields. We first evaluated the survival of each host in the soil collected from each field. Later, the presence of the galls characteristic of clubroot disease was assessed visually and under the stereo microscope Stemi 305 (ZEISS, USA). To confirm the presence of the pathogen, symptomatic plants grown in the soil collected from Tlaxcala and Puebla were selected randomly and resting spores were extracted using a sucrose-based method as previously described (Pérez-López et al. 2021) and observed under the VE-M5D Digital Microscope (VELAB, USA). To fulfill Koch postulates, 1×10^7^ resting spores were used to inoculate the soil where cabbage seeds were germinated and grown.

Total DNA was extracted from 1 g of dry soil from each sample and from all symptomatic roots as previously described (Adame et al. 2016). DNA was diluted in distilled and sterile water and 2-4 ng of DNA was used in 25 μl PCR assays to amplify *P. brassicae* partial 18S rRNA-encoding gene (GenBank accession no. AF231027) with primers TC1F/TC1R and PbF/PbR as previously described (Cao et al. 2007; Wallenhammar et al. 2012). *P. brassicae* DNA was revealed by the presence of ~500 bp amplicons for primers TC1F/TC1R and/or a ~100 bp fragment with primers PbF/PbR, when observed in an ethidium bromide-stained 1 % agarose gel using an ultraviolet transiluminator. Amplicons were purified using QIAquick® PCR Purification Kit (QIAGEN, Mexico) and directly sequenced using the amplification primers.

### Phylogenetic analysis

The partial 18S rRNA-encoding gene sequences generated in this study were assembled using the Staden package (Bonfield and Whitwham 2010) and compared with reference sequences from GenBank using BLAST (http://www.ncbi.nlm.nih.gov). Phylogenetic analysis was conducted using the Maximum Parsimony method with MEGA 6 (Tamura et al. 2013), and bootstrapping 1000 times to estimate stability. *Ligniera* sp. (isolate F69) (AJ010425) was used as outgroup to root the tree.

### Statistical analysis

Survival of hosts by field and control groups were compared using a two-way ANOVA followed by Tukey’s HSD post hoc test was used to compare the mean of multiple groups. All statistical analysis were performed using STATISTICA v.10 (https://statistica.software.informer.com/10.0).

## RESULTS

### Identification of areas putatively infected by the clubroot pathogen

During the interviews with government representatives in Puebla and leaders of grower’ associations in Tlaxcala, we were able to identify several broccoli producers San Pedro Cholula (SPC), Puebla, and Lázaro Cárdenas (LC), Tlaxcala. In conversation with growers, we learned that most of them have seen before clubroot-related symptoms in their fields, and that in several fields the yield has been declining so drastically that growing broccoli or other brassicas crops is not longer profitable. All these elements were pointing to the probable presence of *P. brassicae* in those fields. Interestingly, we also found that in 9 of the fields, representing more than 64% of the fields sampled, are no longer used to grow brassicas (Fig. 1).

### Detection and characterization of the clubroot pathogen in Mexico

Sixty days after transplantation, the four hosts used in this study were evaluated to determine how many survived in the soil collected in every field. Survival was significatively reduced for all the hosts in the soil from the fields collected in San Pedro Cholula, Puebla, ranging from close to 40% survival in plants grown in soil collected from SPC2 to no plants surviving when grown in the soil collected in SPC7 (Fig. 2A, Table 1). In Lázaro Cárdenas, 50-60% of the plants survived 60 dpt, registering the lower survival in LC5 (Fig. 2B, Table 1).

**Fig. 2.**
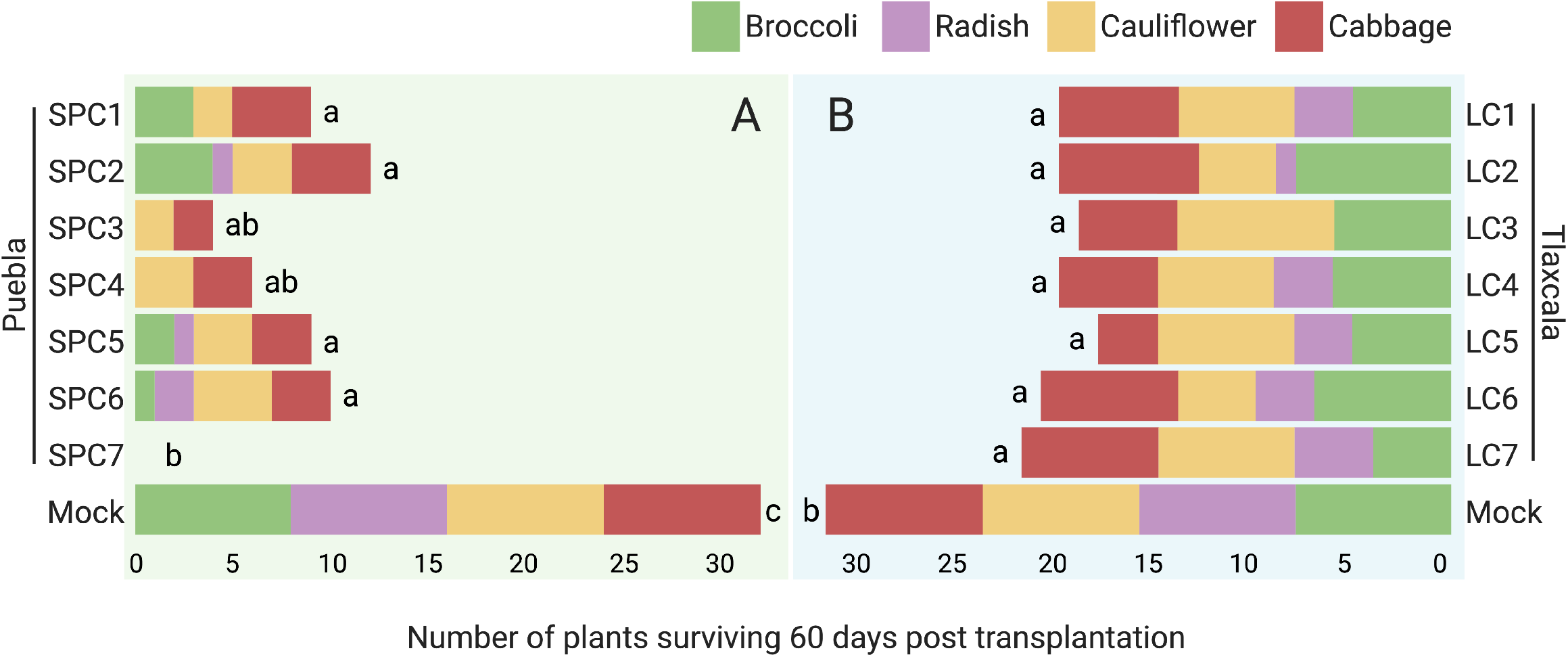
Survival of the four hosts used as bait 60 days post transplantation to the soil collected from the fields evaluated in the study. **A**. Fields from San Pedro Cholula, Puebla. **B**. Fields from Lazaro Cardenas, Tlaxcala. The survival in every field in both geographic locations was significatively lower than in the mock inoculated plants. Identical letters indicate that there is no significative differences among the treatments with *p* ≤ 0.05.

**Table 1.**
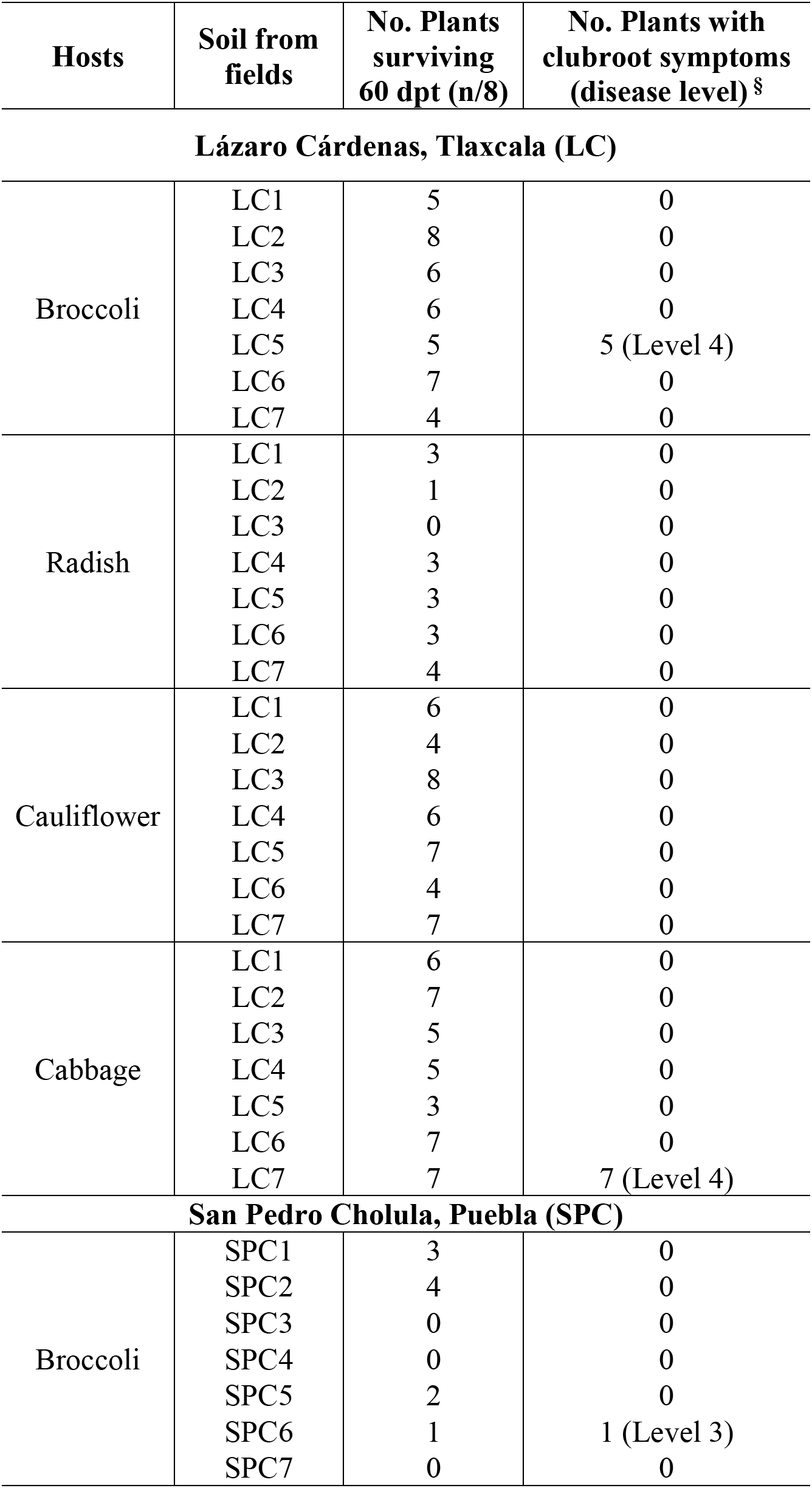
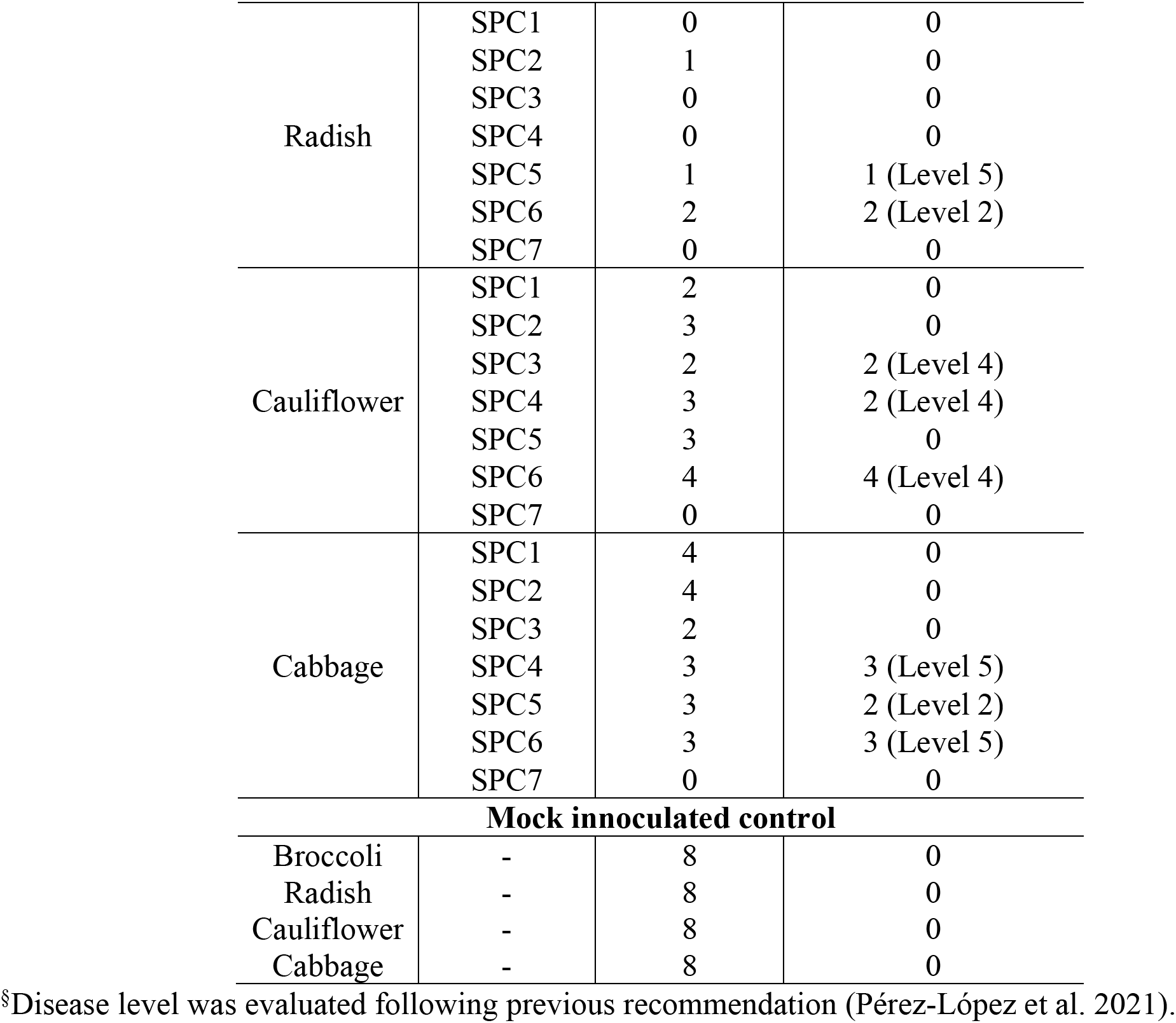
Survival of brassicas hosts grown in the soil collected from Tlaxcala and Puebla and clubroot symptomatology.

After evaluating plants survival, a parameter that could indicate the presence of pathogens in the soil collected, we evaluated the presence of clubroot-characteristic galls in the roots. The first thing we noticed is that the development was seriously affected in all remaining plants at 60 dpt (Fig. 3A). Later, during the root evaluation we were able to identify galls in all of them, rating the disease from level 2 to level 5 (Fig. 3A, Table 1). In Tlaxcala, we only detected symptoms in broccoli plants grown in the soil collected from LC5, and in cabbage grown in the soil from LC7. In Puebla, the region with the lower survival, the plants grown in the soil collected from four out of seven fields showed clubroot symptomatology (Table 1). Overall, of 480 interactions established, 206 were successful in terms of survival 60 dpt but only 32 resulted in plants showing clubroot-related symptomatology, in six of 14 fields sampled (Table 1).

**Fig. 3.**
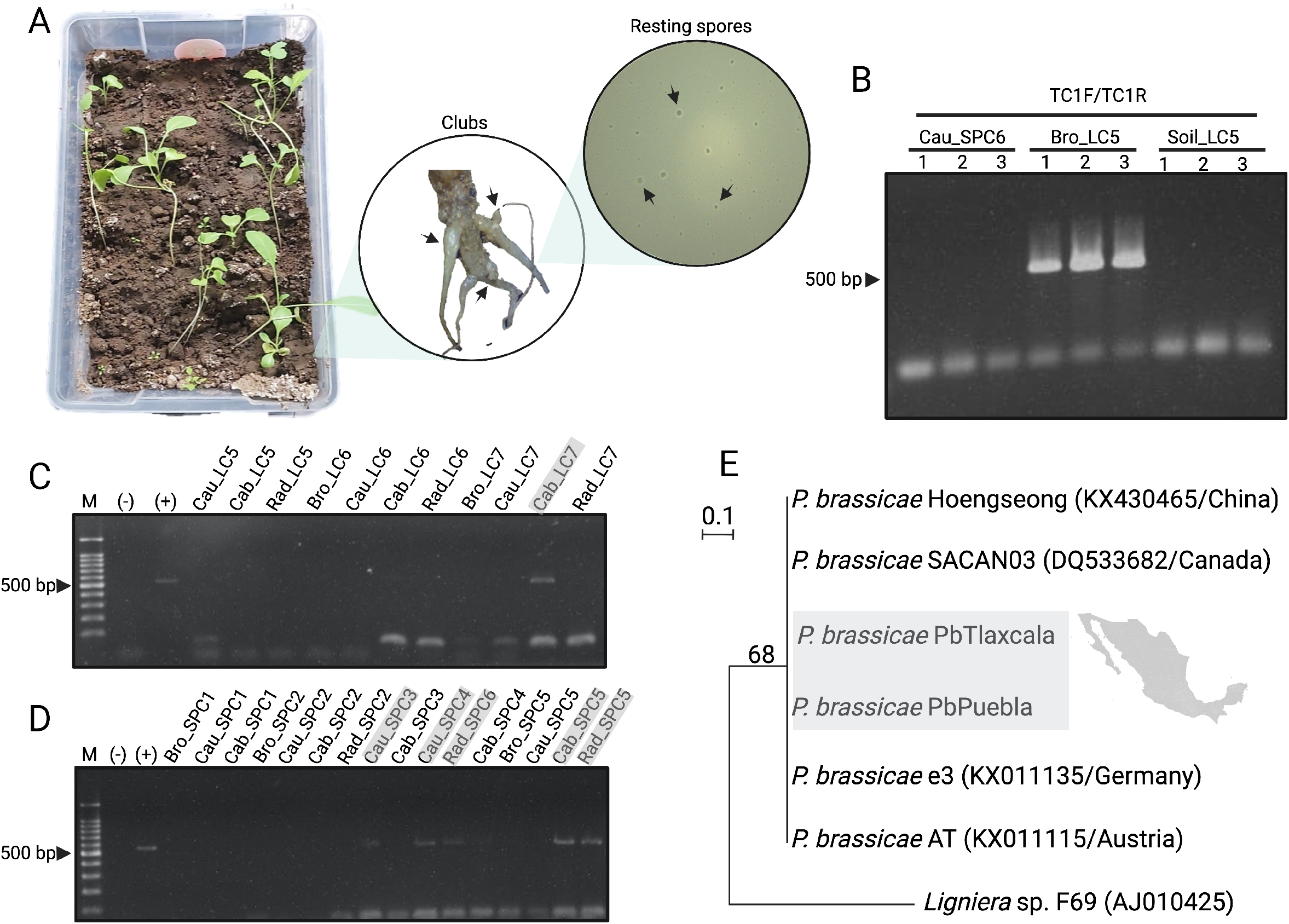
Detection of *Plasmodiophora brassicae* in Mexico. **A**. Brassicas baits 60 dpt, clubroot symptomatology observed in infected hosts and spores isolated from galls. **B**. PCR amplification of the partial *P. brassicae* 18S rRNA-gene using as template Bro_LC5, but not from the soil from the same field, where 1, 2, and 3 refers to 2, 3, and 4 ng od DNA used as template in the PCR assays. The extended version of the figure is Fig. S1. **C**. PCR amplification of the partial *P. brassicae* 18S rRNA-gene using as template symptomatic plants grown in soil from Lazaro Cárdenas, Tlaxcala. **D**. PCR amplification of the partial *P. brassicae* 18S rRNA-gene using as template symptomatic plants grown in soil from San Pedro Cholula, Puebla. The extended version of panels C and D is presented in Fig. S2. **E**. Phylogenetic tree of *P. brassicae* 18S rRNA-gene using *Ligniera* sp. (isolate F69) (AJ010425) as outgroup and bootstrapped 1000 times.

Confirmation of the presence of *P. brassicae* was obtained after the extraction of resting spores from symptomatic roots of randomly selected plants. From all of them we were able to observe resting spores showing their characteristic rounded morphology (Fig. 3A). The spores were used to inoculate cabbage plants and in 90% of the plants inoculated we were able to observe clubroot-related symptoms. These result were confirmed through PCR assays targeting *P. brassicae* genomic DNA (Fig. 3B, Fig. S1). To test the two set of primers we first used the DNA extracted from cauliflower grown in SPC6 soil (Cau_SPC6), broccoli grown in LC5 soil (Bro_LC5), and soil collected in LC5 (Soil_LC5). Using primers PbF/PbR we were not able to observe the expected ~100 bp amplicon (Fig. S1), while using primers pair TC1F/TC1R, the expected ~500 bp amplicon was only observed when genomic DNA from Bro_LC5 was used (Fig. 3, Fig. S1). Moving forward we decided to use the DNA from the 32 symptomatic plants to confirm the presence of the clubroot pathogen in the fields in Tlaxcala and Puebla. All symptomatic plants grown in the soil collected from LC5 (Broc_ LC5) and LC7 (Cab_ LC7) in Tlaxcala were positive for *P. brassicae* by PCR (Fig. 3C, Fig. S2). In Puebla, all symptomatic plants grown in the soil collected from SPC3 (Cau_ SPC3), SPC5 (Rad_ SPC5 and Cab_ SPC5), and SPC6 (Bro_ SPC6, Rad_ SPC6, Cau_ SPC6, and Cab_ SPC6) were positive for the clubroot pathogen (Fig. 3D, Fig. S2). Curiously, only symptomatic cauliflower plants grown in the soil collected from SPC4 (Cau_ SPC4) were positive for the presence of *P. brassicae* by PCR, while symptomatic cabbage plants (Cab_ SPC4) were negative (Fig. 3D).

To confirm that the amplicons observed were indeed *P. brassicae*, the purified PCR product obtained using total DNA from Bro_LC5 and Cab_SPC5 plants was purified and sequenced. All the sequences were 100% identical among them and 100% identical to the 18S rRNA-encoding gene of *P. brassicae* isolates from China (KX430465), Canada (DQ533682), Germany (KX011135), and Austria (KX011115). Blast results were supported by the phylogenetic tree obtained by maximum parsimony analysis of the 18S rRNA-encoding gene of *P. brassicae Pb*Tlaxcala (Genbank accession no. OL461232) and *Pb*Puebla (Genbank accession no. OL461231) (Fig. 3E).

## DISCUSSION

Clubroot has been a constant threat to brassicas production for more than 100 years in The Americas (Dixon, 2009). The growing market and interest for brassicas vegetables and oilseed crops has expanded to Central and South America and with it the needs to know more about the distribution of plant disease that could put in danger the economic profit linked to those crops. In this study, on the heels of true detective work, was possible to confirm for the first time the presence of *Plasmodiophora brassicae* in Mexico. Our results provide a comprehensive and detailed identification of the clubroot pathogen in soil collected from two of the main broccoli producer areas in Eastern Mexico, findings of great impact for the future management of the disease and the pathogen.

The first challenge that we faced was to find the fields to perform sampling. After several meetings and discussions with growers’ associations and government representatives from the states of Puebla and Tlaxcala we were able to identify several growers that have previously seen clubroot symptoms in their fields. Although anecdotical, the growers mentioned that they have had to stop growing broccoli in several of their fields because plants development has been seriously affected lately, and subsequently yield and profit. We were then wondering if these issues could be related to the presence of *P. brassicae* in the area, motivating the collection of soil. For the purpose of this study, fourteen geographically distanced broccoli fields, and a total of 70 soil samples were collected.

In this study we had the fortuitous opportunity to assess to three different group of fields: fields in production (LC6, SPC1, SPC3, SPC4, and SPC7), fields without brassicas from 2 months to 1 year (LC3, LC5, SPC2, SPC5, and SPC6), and fields without brassicas for 2 years or more (LC1, LC2, LC4, and LC7). Plants grown in the soil collected from SPC fields in Puebla had the lower survival rate compared to the survival registered in the soil collected from Tlaxcala. This was consistent with the presence of clubroot symptomatology, registering a higher number of symptomatic plants among those grown in the soil collected in Puebla. Another exciting finding is that 4 of the 6 fields where the clubroot pathogen was detected were fields where growers had to stop growing broccoli or other brassicas due to the difficulties to obtain the expected yield and profit. Those where LC5, SPC5, and SPC6 ranging from 2 months to 1 year without growing brassicas on them, and LC7 without brassicas for 2 years or more.

Something important to mention is that the strategy followed for us to investigate the presence of *P. brassicae* could have failed in detecting the pathogen in some fields, mainly because we did not use the DNA extracted from the soil as template for the PCR assays. The main reason of this decision was the first test confirming that although Soil_LC5 was negative for *P. brassicae*, Bro_LC5 was positive, evidencing that plant DNA was a more reliable than soil DNA. The detection of soil-borne pathogens using soil samples is a big challenge (Wallenhammar et al. 2012). The high amount of PCR inhibitors in the soil can lead to high rate of false negatives, and it is challenging starting from environmental samples with unknown amount of inoculum to “catch” the pathogen (Wen et al. 2020). The use of bait crops has been suggested and evaluated to trap spores in Canadian fields, showing that they can successfully be infected by the clubroot pathogen even when the number of resting spores is low (Ahmed et al. 2011). The strategy followed, although might allowed to miss the presence of the pathogen in some fields, did not changed the main goal of the study, the confirmation of the presence of the clubroot pathogen in Mexico.

Another exciting finding was the connection between plants survival and clubroot symptomatology. Although we did not perform statistical analysis to support the correlation among hosts survival, symptomatology, and presence of the clubroot pathogen, we observed a clear trending indicating that in the soil where the survival of the hosts evaluated was lower, the number of plants affected by clubroot disease was higher. These findings are clearly pointing to the presence of the clubroot pathogen as the main reason why the profitability of broccoli and other brassica vegetables has drastically reduced in some areas of the states of Puebla and Tlaxcala.

Clubroot management has proven to be very challenging, and it has relied mainly in stopping the introduction of the pathogen into clean fields (Botero et al. 2019; Dolatabadian et al. 2021). However, how to avoid the spreading of a pathogen you don’t know you have in your country or even geographic area? Studies like this one developed by us are key for diseases management, especially for those devastating like clubroot, most likely already affecting the Mexican economy although no official research has taking place. To contribute to the management of the disease we have added, with the agreement of the growers, the location of those fields where the presence of the clubroot pathogen was confirmed to the interactive online tool ClubrootTracker (http://clubroottracker.ca) (Muirhead et al. 2020) (Fig. 4). Now, any grower in the area can have access to the current confirmed distribution of the clubroot pathogen in Tlaxcala and Puebla and take measures to avoid the spreading of the pathogen. To ensure the utility of the new data for Mexican growers the message that opens once the location has been clicked is in English and Spanish (Fig. 4).

**Fig. 4.**
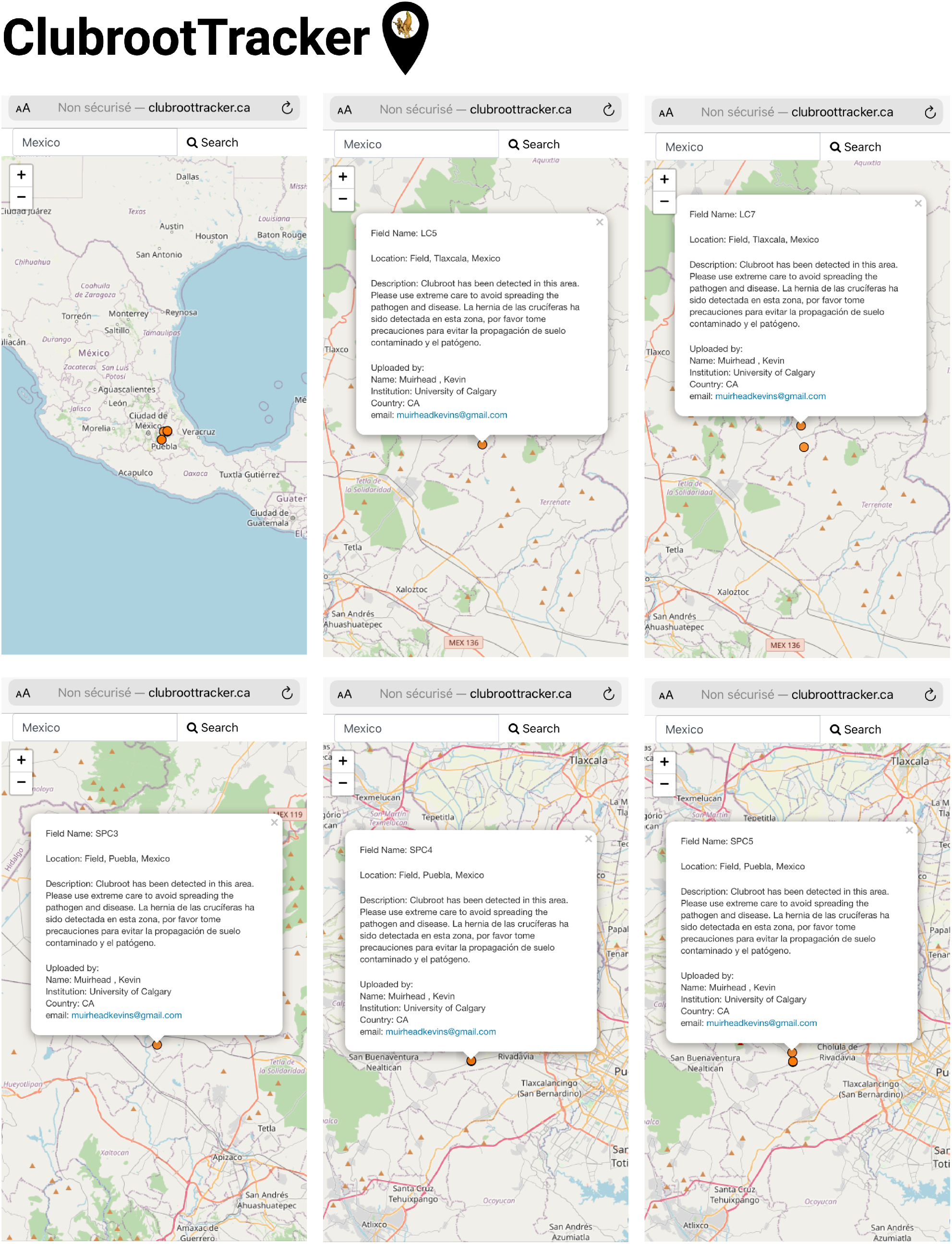
ClubrootTracker display of the fields confirmed positive for *P. brassicae* in Puebla and Tlaxcala. The information added to each positive location is also display in the figure.

In conclusions, the work presented here is the first molecular identification of *P. brassicae* in Mexico, making official the presence of the clubroot pathogen in all North America. This study has opened the door to a new area of research in Mexico focusing on the distribution of clubroot disease, the best management strategies depending on the local conditions, and the search for resistant varieties to mitigate the damage caused by the disease in areas already affected. Our results are also the first step to fully characterize for the first time the genome of a *P. brassicae* isolate from Latin America. This will contribute to finally identify for example, the core effector repertoire of the pathogen and characterize conserved mechanisms used by the clubroot pathogen to infect the susceptible host. We are looking forward to seeing all the exciting and new research flourishing from our results.

## ACKNOWLEDGES

We would like to thanks to the authorities of Puebla and Tlaxcala for the support. Legnara Padrón-Rodríguez thanks the Posgrado en Ciencias Agropecuarias, Facultad de Ciencias Agrícolas, Universidad Veracruzana and CONACYT for the PhD scholarship (CVU: 894229). Authors thanks also Kevin Muirhead for updating ClubrootTracker with the location of the clubroot distribution in Mexico.

---

**Fig. S1.**
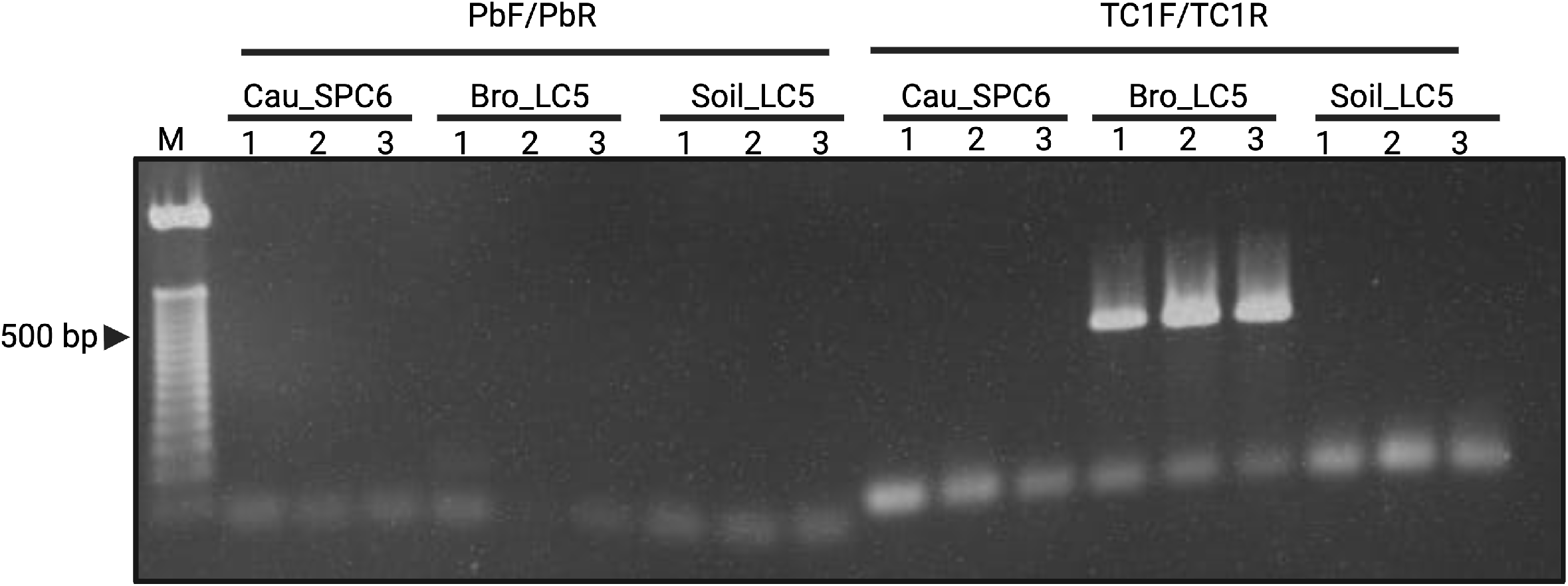
Extended figure 3B. Detection of *Plasmodiophora brassicae* using plant tissue and soil with primer pairs PbF/PbR and TC1F/TC1R. No clear band was observed with primers PbF/PbR, while with primers TC1F/TC1R the amplicon of the expected size was observed when total DNA from broccoli (Bro) in LC5.

**Fig. S2.**
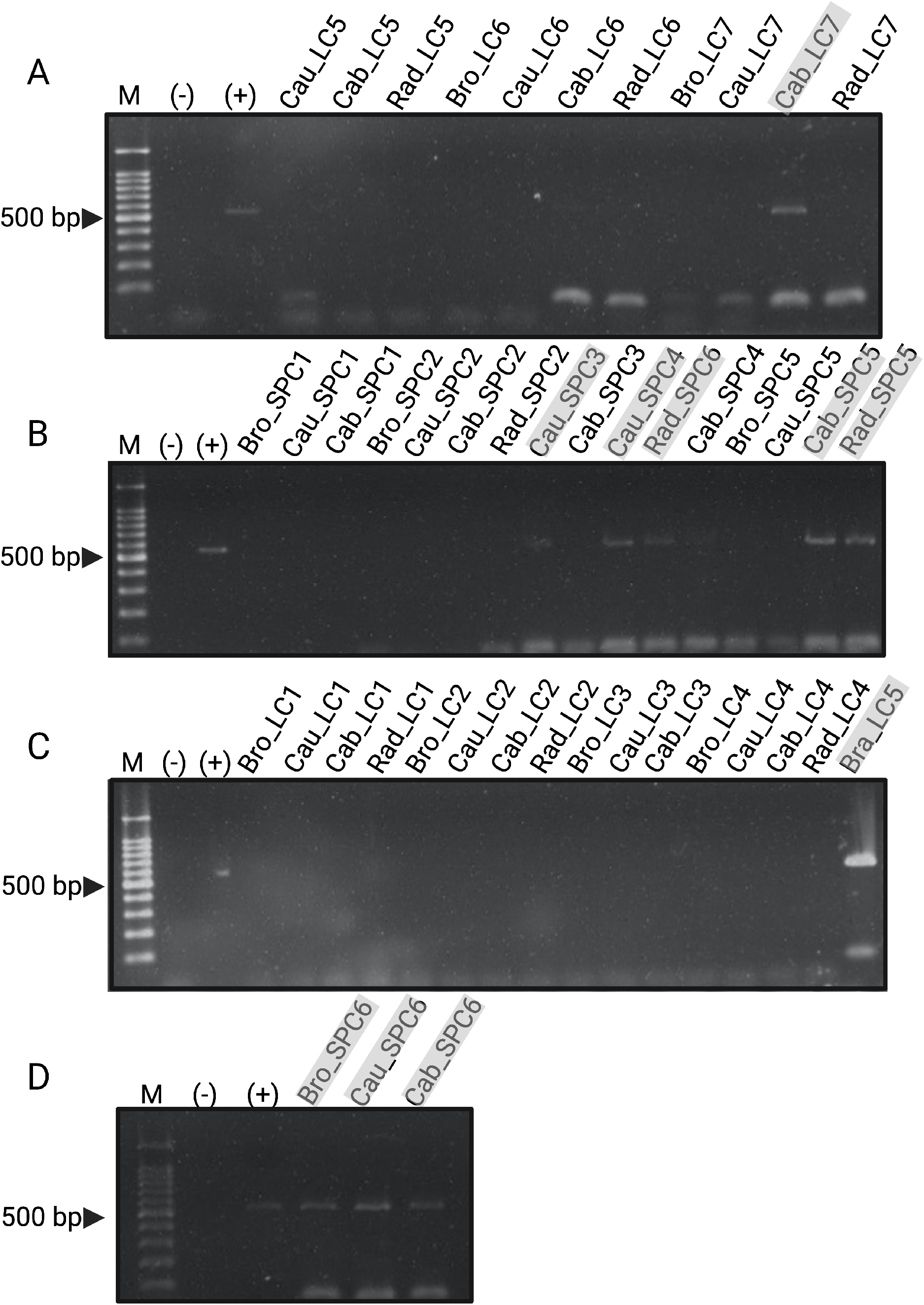
Extended figure 3C-D. Detection of *Plasmodiophora brassicae* using plant tissue of plants grown in soil collected in Tlaxcala fields (LC), and Puebla fields (SPC) with primers TC1F/TC1R. Cau stands for cauliflower, Bro, broccoli, Rad, radish, and Cab, cabbage.

